# An efficient one-step rRNA depletion method for RNA sequencing in non-model organisms

**DOI:** 10.1101/2025.08.15.670489

**Authors:** Muhammad Suleman Qasim, L. Peter Sarin

## Abstract

RNA sequencing (RNA-seq) has revolutionized global transcriptomic analysis, ribosome footprinting, and polysome profiling, providing a wealth of data. Importantly, many RNA-based omics approaches typically involve either the removal of ribosomal RNA (rRNA) or selection of messenger RNA (mRNA) prior to sequencing, thereby enriching reads that map to the translationally active part of the transcriptome. Prokaryotic mRNA differs from eukaryotic mRNA in that it lacks the 3’ polyadenylated tail, which excludes the use of poly(A)-based selection methods. While commercial rRNA depletion products exist for a growing number of prokaryotes, their proprietary nature and potential inefficiency with non-model organisms are factors that may limit broad-scale application. To mitigate this issue, we designed DepStep, a consolidated workflow for one-step rRNA depletion using species-specific biotinylated antisense probes for selective hybridization and removal of the target rRNA molecules. As a proof-of-concept, RNA-seq libraries of the psychrophilic gram-negative bacterium *Shewanella glacialimarina* TZS-4_T_ were prepared using both DepStep and a commercial rRNA depletion kit for gram-negative bacteria, to which DepStep was benchmarked. DepStep compares favorably to the commercial depletion kit; it efficiently removes >98.6% of the rRNA content, and a slight increase in total read counts aligning to the coding sequences (CDS) was observed. Importantly, DepStep’s cost-per-sample is three times lower than the commercial kit, establishing DepStep as a simple yet cost-effective alternative to commercial solutions.

## INTRODUCTION

RNA sequencing (RNA-seq) has become a pivotal tool for generating comprehensive transcriptional profiles and uncovering mechanistic insights into various biological processes (Wang et al. 2009; Ozsolak and Milos 2011). However, a significant challenge in RNA-seq is the overwhelming presence of ribosomal RNA (rRNA), which constitutes up to 90% of the total RNA in both prokaryotes and eukaryotes (Deng et al. 2022). This abundance of rRNA can obscure the detection of messenger RNA (mRNA) during sequencing, thereby skewing the transcriptomic data and limiting the insights that can be derived from protein-coding sequences. In eukaryotes, this issue is often circumvented by poly(A) enrichment, a technique that selectively isolates polyadenylated mRNA for sequencing (Mortazavi et al. 2008). However, this approach is not applicable to prokaryotes, as most prokaryotic mRNAs lack polyadenylation. Hence, rRNA must be effectively removed from the total RNA to allow accurate quantification of mRNA transcripts.

To achieve this, the preferred option is to rely on commercial rRNA depletion methods, which are available from numerous vendors, particularly when working with established model organisms. On the other hand, commercial kits may be less efficient with non-model organisms, as their depletion strategies might not be as effective in removing rRNA from lesser-studied prokaryotes. Consequently, customized solutions may be needed to remove rRNA, including (i) depletion using rRNA-specific biotinylated probes coupled to streptavidin beads (Thompson et al. 2020; Culviner et al. 2020; Kraus et al. 2019), (ii) targeted degradation of rRNA-DNA hybrids using RNase H treatment (Huang et al. 2020), (iii) specific cDNA synthesis from mRNA using non-random primers that do not prime reverse transcription of rRNA molecules (Armour et al. 2009), and (iv) enrichment of mRNA by blocking the amplification of rRNA (Wangsanuwat et al. 2020). While these methods have been successfully used to enhance the capture of mRNA and enable quantitative analysis of mRNA transcripts, they are often laborious to implement, requiring rigorous testing before optimal performance is achieved.

Drawing from these strategies, here we present a simplified one-step rRNA depletion method – DepStep – that utilizes biotinylated antisense rRNA probes linked to streptavidin-coated magnetic beads. Using the non-model bacterium *Shewanella glacialimarina* TZS-4_T_ as proof-of-concept, we designed species-specific rRNA antisense probes and systematically optimized the depletion conditions, removing >98.6% of rRNA molecules without biasing the mRNA and non-coding RNA contents. To evaluate the effectiveness of our approach, we compared our in-house method to a commercially available rRNA depletion kit that is compatible with our non-model organism. We demonstrate that DepStep offers similar depletion as the commercial kit and mRNA reads from both methods showed a positive Pearson correlation (P = 0.84). Importantly, we noticed that uniform probe coverage throughout the rRNA molecule is critical for its effective removal and maintaining a maximum distance of less than 200 nt between adjacent probes resulted in optimal depletion efficiency. Moreover, DepStep can be easily adapted to generate biotinylated probes for other organisms, facilitating studies on transcriptional regulation across different species. By employing DepStep, we enhanced the detection of mRNA transcripts and increased the overall reads coverage across the CDS region in *S. glacialimarina*. Consequently, DepStep provides a consolidated workflow for generating custom probes, offering an efficient and cost-effective solution for rRNA removal.

## RESULTS

### Uniform probe coverage is critical for efficient rRNA removal

The *S. glacialimarina* genome contains 22 rRNA gene copies, including 7 copies of the 23S rRNA gene, 7 copies of the 16S rRNA gene, and 8 copies of the 5S rRNA gene. To deplete bacterial rRNA originating from both the large and the small ribosomal subunits, a set of antisense oligonucleotides were designed to target each rRNA molecule. First, biotin-tagged antisense oligonucleotides were pooled, then hybridized to rRNA and later crosslinked to streptavidin beads. This captures the DNA probes along with the bound rRNA molecules, effectively removing the rRNA from the total RNA sample (Fig. 1A). In DepStep, probe lengths were maintained between 26-32 nt to ensure a minimum ΔG value for hybridization and a melting temperature >70 °C, which allows the RNA-DNA hybrids to remain stable during the streptavidin capture step. The potential probe candidates were generated with Oligostan.R (Tsanov et al. 2016) using default parameters. A total of 83 candidate sequences were generated: 54 probes targeting 23S rRNA, 27 probes targeting 16S rRNA, and 2 probes targeting 5S rRNA. Next, candidate sequences were screened for hairpin/dimer formation and *in silico* cross-hybridization analysis using BLASTn from NCBI (Altschul et al. 1990) and BURST v0.99.8 (Al-Ghalith and Knights 2020), excluding probes with significant homology to the CDS region. The abovementioned steps are outlined in the DepStep rRNA depletion workflow (Supplemental Methods). Following the implementation of quality control measures, the final probe library contained 35 probes; 20 targeting 23S rRNA, 13 targeting 16S rRNA, and 2 targeting 5S rRNA (Supplemental Table 1).

**Figure 1.**
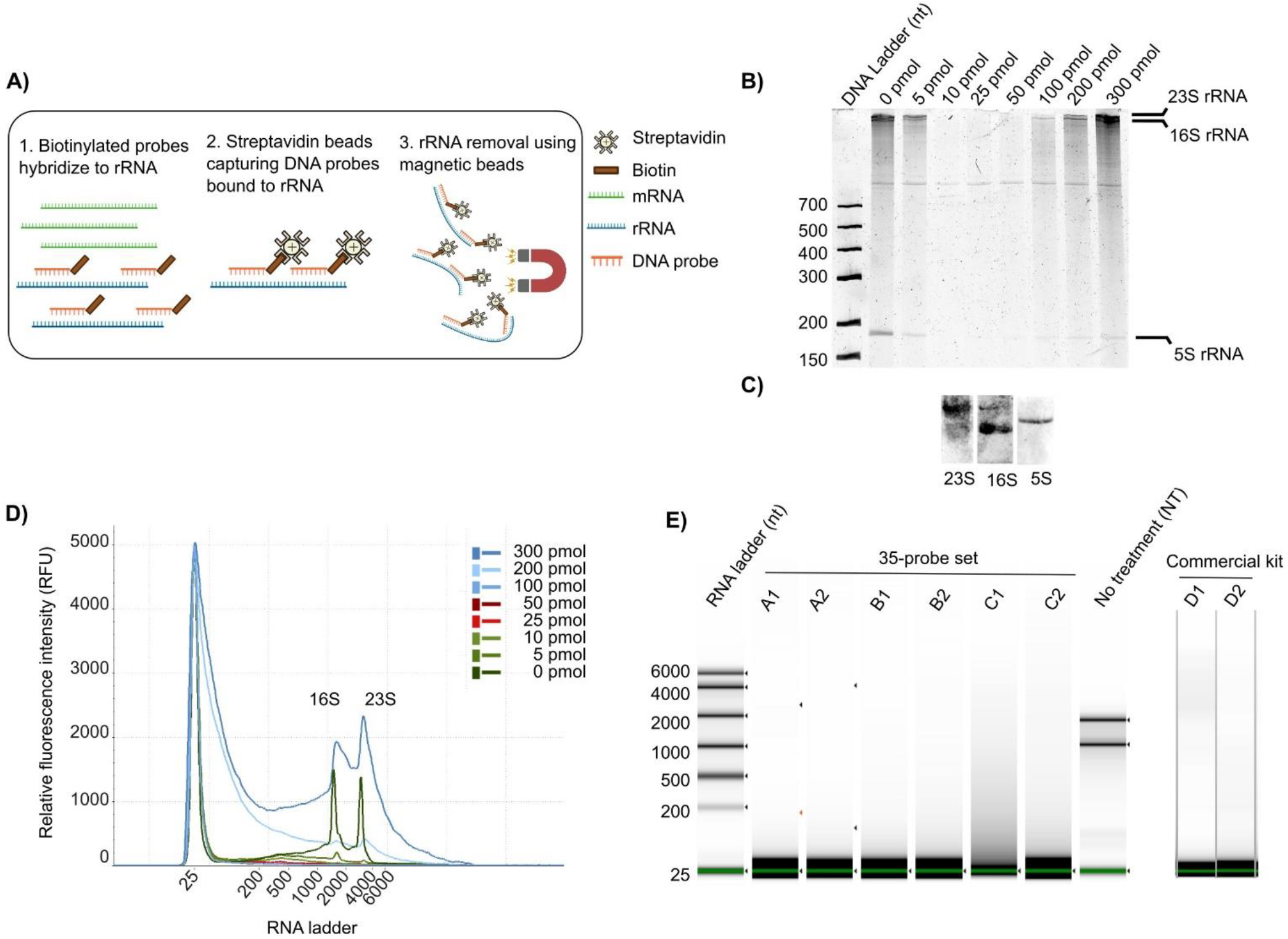
DepStep rRNA depletion method using biotinylated antisense probes. A) Schematic representation of the rRNA depletion method. B) Representative 6% urea polyacrylamide gel electrophoresis of *S. glacialimarina* TZS-4_T_ total RNA samples following rRNA depletion using 35-probe set mix with increasing probe amount (from 0 pmol to 300 pmol). Size labels for 23S, 16S and 5S rRNA are shown on the right side of the gel, whereas DNA molecular weight markers are shown on the left side of the gel, respectively. C) Representative northern blot of 35-probe set against *S. glacialimarina* 23S, 16S and 5S rRNA. D,E) Automated electrophoresis chromatograms of (D) rRNA depletion samples after depletion with increasing concentration of 35-probe set mix (from 0 pmol to 300 pmol) and (E) rRNA-depleted samples after 35-probe set mix depletion (50 pmol) and the commercial depletion kit. For each sample, the alphabetical label denotes an independent biological sample whereas numerical labels denote technical replicates of each biological sample. Fig 1A was created using Biorender.

To estimate the number of probes needed for efficient rRNA depletion with DepStep, we generated two probe sets from the final probe library; a limited 22-probe set featuring the most optimal probe designs (10 probes for 23S rRNA, 10 probes for 16S rRNA, and 2 probes for 5S rRNA), and the complete 35-probe set that maintains an even distribution of probes across all rRNAs (one probe per ∼150 nt, maximum distance between adjacent probes not exceeding 200 nt) (Supplemental Fig. 1A; 35-probe set). Following rRNA depletion from 1 µg of total RNA and subsequent automated electrophoresis chromatogram analysis, we observed that the 35-probe set effectively removed rRNA, leaving no visible rRNA bands in the samples. Conversely, the limited 22-probe set did not achieve a similar level of rRNA depletion as two faint but clearly distinguishable bands were present in all samples (Supplemental Fig. 1B). Since the bands were absent in the 35-probe set treated samples, we hypothesized that they might represent portions of the 23S rRNA molecule that were not targeted by the 22-probe set (Supplemental Fig. 1A; 22-probe set). Hence, we concluded that uniform coverage of the probe, with an approximate spacing of no more than 200 nt between adjacent probes, is important to ensure efficient removal of all rRNA and prevent rRNA fragmentation.

### Optimal probe amount ensures efficient rRNA removal

Accurately estimating the probe amount relative to the input total RNA is critical for optimizing the depletion efficiency and reducing cost. If too much probe is applied, the rRNA substrates become saturated, resulting in excess unbound probes that interfere with and reduce the efficiency of streptavidin capture of hybridized (rRNA bound) probes. Hence, these unbound probes must be efficiently removed, necessitating a higher amount of costly streptavidin beads to purge the residual probes from the depletion reaction mix. Meanwhile, a significant portion of the streptavidin beads remains linked to unhybridized probes. To determine the optimal probe concentration, we tested the depletion of rRNA from 1 µg of total RNA using probe amounts ranging from 5 pmol to 300 pmol of 35-probe set mix. This approach enabled us to explore the upper and lower limits of the probe quantity required for rRNA depletion. Gel electrophoresis analysis for the 35-probe set mix indicated that a probe concentration from 10 – 50 pmol effectively removed the majority of 23S, 16S, and 5S rRNA (Fig. 1B). With Northern blots we confirmed that the 35-probe set for 23S, 16S and 5S sequences was able to bind to the target rRNA sequences (Fig. 1C). Similarly, the automated electrophoresis chromatogram of the depleted rRNA samples showed a progressive decrease in the amount of 23S and 16S rRNA by up to 98.6% when using 10 pmol, 25 pmol, and 50 pmol probe amount, respectively (Fig. 1D). For the final RNA sequencing library preparation, we proceeded with 50 pmol of 35-probe mix per µg of total RNA, and the depletion efficiency was assessed by electrophoresis chromatogram analysis (Fig. 1E). This analysis confirmed that rRNA depletion with 50 pmol of 35-probe mix was successful with no detectable rRNA bands appearing in the electrophoresis chromatogram. Hence, to assess the rRNA depletion potential of DepStep, we acquired several commercially available rRNA depletion kits against gram-negative bacteria for comparison. However, only RiboCop rRNA depletion kit (Lexogen; see Materials and Methods) successfully depleted *S. glacialimarina* rRNA, and this kit was subsequently selected as the benchmark for DepStep. Depletion reactions were performed in triplicate with each biological sample having two technical replicates for both DepStep and the commercial kit. Importantly, DepStep and the commercial kit yielded comparable results in rRNA removal efficiency (Fig. 1E). Therefore, our results demonstrate that DepStep, when implemented with optimized probe amount and strategically selected target regions, effectively eliminates the majority of rRNA for the total RNA sample.

### DepStep rRNA depletion increases CDS reads to >80% of the total read count

Sequencing libraries were prepared using rRNA-depleted samples obtained from DepStep (22 and 35-probe sets) and the commercial kit, including libraries from total RNA with no treatment (NT) samples as a control. After sequencing, the reads were mapped back to the reference genome and the sum of reads aligning to 23S, 16S, 5S rRNA and the CDS regions were compared between each group of samples. DepStep depletion using the 35-probe set (DepStep-35P) proved efficient, as reads mapping to rRNA accounted for <20% of the total read count (Fig. 2A-C), except for sample 35P-A1 (attributed to a technical error) (Supplemental Fig. 1C). However, the 22-probe set (DepStep-22P) was largely inefficient in removing rRNA, as most reads aligned to two distinct locations within the 23S rRNA (Fig. 2C). Indeed, these locations were not covered by DepStep-22P, confirming our initial hypothesis that the residual RNA fragments did originate from 23S rRNA fragmentation, which led to parts of 23S rRNA being retained in the final sequencing libraries (Supplemental Fig. 1). By comparison, the commercial depletion kit performed slightly better than DepStep-35P with <10% of the reads mapping to rRNA (Fig. 2A-B). Nonetheless, both depletion methods were successful in increasing the usable CDS read count to >80% of total reads, whereas CDS reads accounted for as little as 1% of total reads in the non-depleted control sample (Fig. 2B). Moreover, read coverage plots also show little to no reads mapping to rRNA for either DepStep-35P or the commercial kit (Fig. 2C). Hence, we also performed a log_2_ fold change analysis of DepStep-35P samples and the commercial kit, comparing both to the non-depleted control. This revealed that both depletion methods reduced rRNA by 8-to 10-fold relative to the control (Fig. 2D). Consequently, DepStep-35P efficiently reduced rRNA from the total RNA sample and the depletion potential is comparable to that of the commercial rRNA depletion kit for gram-negative bacteria.

**Figure 2.**
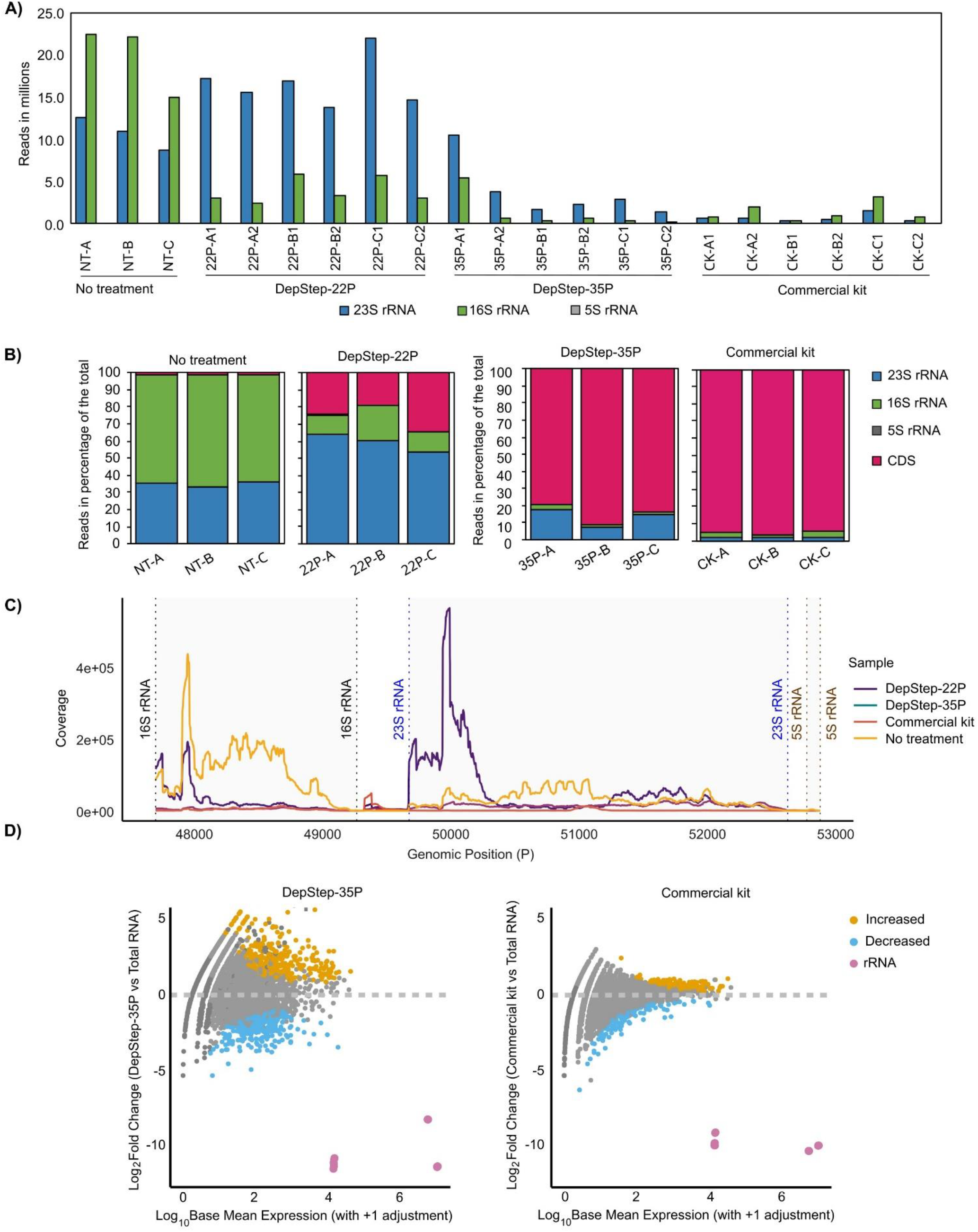
Depletion analysis from DepStep sequencing library preparation of *S. glacialimarina* TZS-4_T_. A) Raw reads count corresponding to *S. glacialimarina* 23S, 16S and 5S rRNA in total RNA with no treatment (NT), DepStep-22P (22P) depletion, DepStep-35P (35P) depletion, and commercial kit (CK). For each sample, the alphabetical label denotes an independent biological sample whereas numerical labels denote technical replicates of each biological sample. B) Bar plots depicting raw read counts (in %) mapping to 23S rRNA, 16S rRNA, 5S rRNA, or CDS in total RNA with no treatment (NT) as well as DepStep-22P (22P), DepStep-35P (35P), and commercial kit (CK) rRNA-depleted samples. C) Coverage plot of reads aligning to 23S, 16S and 5S rRNA sequences in *S. glacialimarina* in DepStep-22P, DepStep-35P, commercial kit and total RNA with no treatment. D) MA plot comparing DepStep-35P (left panel) with the commercial rRNA depletion kit (right panel) with respect to total RNA. The CDS fold change (vs total RNA) for DepStep-35P and the commercial kit are represented as orange (increase) or light blue (decrease) dots, respectively. Magenta dots represent rRNA reads, which are significantly reduced in the rRNA-depleted sequencing libraries.

### CDS reads obtained from DepStep-35P and commercial kit show a strong correlation

Since DepStep-35P and the commercial kit successfully depleted a majority of the rRNA, we expected the relative abundance of mRNA reads to be comparable between both methods. Indeed, the Pearson correlation value of log_10_ transformed and DESeq normalized read counts between DepStep-35P and the commercial kit was 0.82 (Fig. 3A). This represents a positive association and suggests that there is no significant difference between these two depletion approaches. However, it is important to note that the starting material (input total RNA) was prepared separately for each depletion group, which may impact the correlation between the mRNA reads. Therefore, a degree of bias could be accounted due to the different input samples used for each group. Next, we examined the transcript per million (TPM) reads for CDS and focused on CDS associated with known housekeeping genes. Both DepStep-35P and commercial kit samples showed similar TPM trends for each housekeeping gene, albeit the TPM for the DepStep-35P sample was slightly higher than that of the commercial kit (Fig. 3B). Upon closer inspection, we noticed that the DepStep-35P sample had a higher number of CDS with TPM values exceeding 10 and 50, reflecting a general trend of increased reads per CDS (Fig. 3C). Overall, we concluded that the DepStep-35P method efficiently removed rRNA from our sample, enriching mRNA for sequencing. Importantly, DepStep produced comparable results to the commercial rRNA depletion kit with potential improvement in detecting overall CDS expression over the commercial kit.

**Figure 3.**
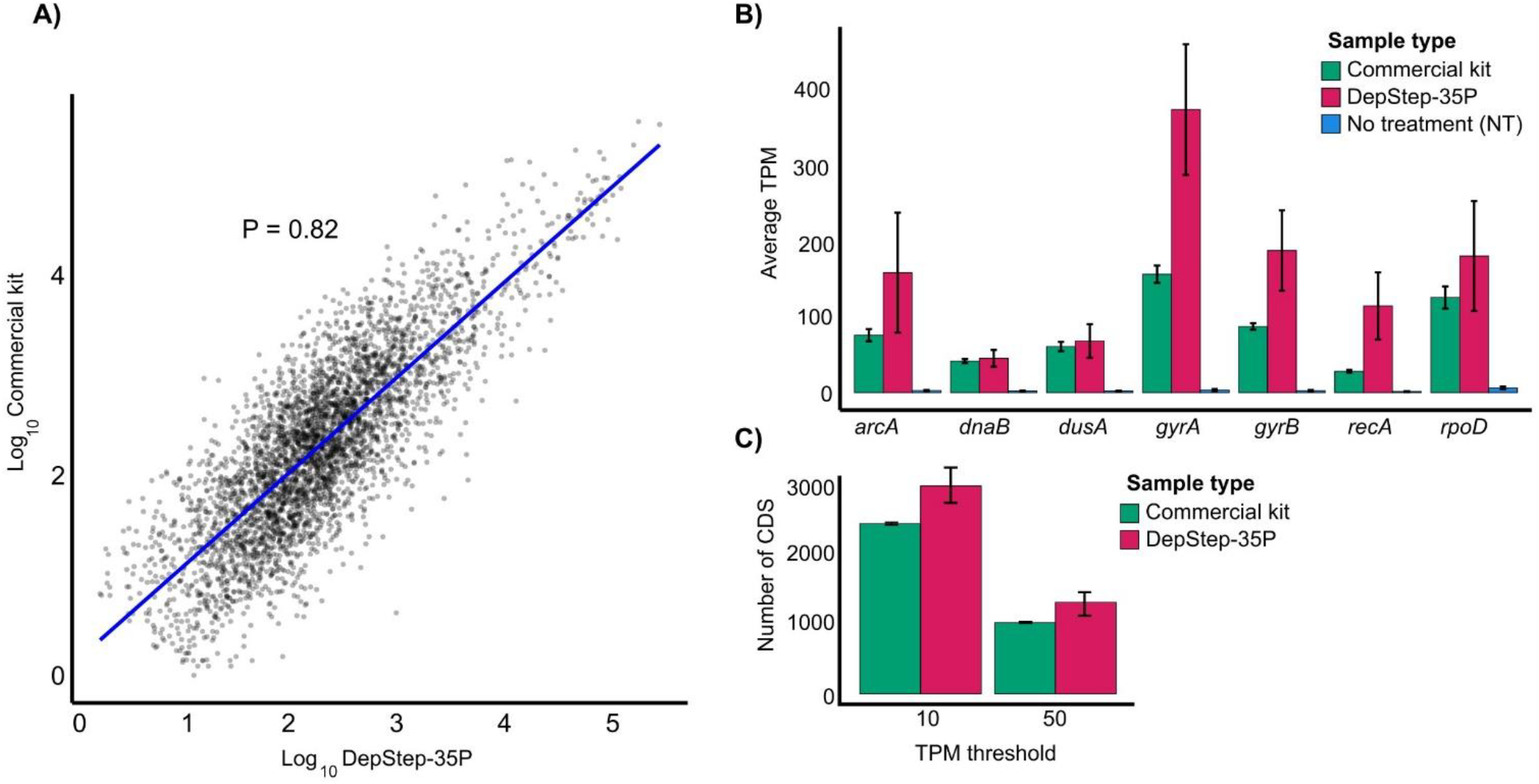
Comparative analysis of coding sequences (CDS) derived from sequencing libraries prepared using DepStep-35P and commercial kit. A) Scatter plot showing Pearson’s correlation between DepStep-35P and commercial kit samples. The RNA levels were estimated by averaging normalized log transformed read counts of three biological replicates for each condition. Lowly expressed genes were removed (log threshold value >0.5). The Pearson’s correlation coefficient is denoted by P. B) Average TPM of common housekeeping genes in rRNA-depleted sample using DepStep-35P and commercial kit vs total RNA. C) CDS count exceeding the threshold value of 10 and 50 for DepStep-35P and the commercial kit, respectively. Error bars represent the standard deviation of three biological replicates.

### DepStep does not lead to off-target binding or introduce other transcriptome biases

To assess for potential off-target binding that could bias transcriptome analysis, we screened the DepStep depletion probe sequences against the *S. glacialimarina* transcriptome using BLASTn, which did not identify any significant off-target effects. Differential expression analysis further confirmed that reads associated with only *S. glacialimarina* rRNA were significantly reduced (5-to 10-fold) after depletion while increasing the TPM of all the other reads in the depleted samples. As for the commercial kit, since the target list of sequences of rRNA was not available, we were unable to directly assess its off-target effects on *S. glacialimarina*. Despite this, we did not observe that reads associated with CDS or TPM would have been adversely affected by the commercial rRNA depletion kit. Therefore, it can be concluded that both methods exhibit minimal off-target effects.

### DepStep rRNA depletion has a favorable per sample cost compared to commercial alternatives

The primary cost in DepStep is attributed to the use of species-specific biotinylated probes and streptavidin beads. Combined, the total material cost is ∼20 €/reaction, i.e. significantly less than 62 €/reaction for the commercial alternative used in this study. Although DepStep has a favorable cost-to-performance ratio, it is worth noting that it may be less labor-intense to opt for a commercial kit, provided the product is compatible for use with the organism-of-study. All materials were purchased from local vendors, and the reported costs were calculated at time of experimentation.

## DISCUSSION

DepStep is a rapid and simple to use single-step method for targeted removal of rRNA – a critical step in RNA-seq that enables a wider coverage of reads across CDS regions (Fig. 1A). To generate this customized solution, we optimized (i) the probe amount, (ii) the number of probes per nt of the target sequence, and (iii) the amount of starting material (total RNA). First, to obtain an efficient depletion of rRNA, the probe amount must be adjusted relative to the starting material (Fig. 1B). Indeed, the depletion process becomes less efficient if the probe amount is too high, as streptavidin beads primarily sequester unbound probes rather than rRNA-probe complexes. This can be mitigated by adding more streptavidin beads, which increases the per-sample cost of rRNA depletion, or preferably by reducing the probe amount. Next, we determined that for optimal rRNA depletion, an average probe density of one probe per 150 nt rRNA is necessary while maintaining a maximum distance of 200 nt between adjacent probes. This uniform distribution proved critical, as a reduced probe coverage resulted in rRNA fragmentation and significantly higher percentage of sequencing reads mapping to rRNA. Therefore, we strongly recommend performing a quality control step after DepStep depletion to validate the extent of rRNA removal. Our analysis showed that DepStep was able to remove a majority of all rRNA, enriching for mRNA transcripts (Fig. 2A,B). Although our results indicate that the commercial kit was more efficient in depleting rRNA, DepStep achieved a CDS count to ∼80% of the total reads, which enabled reliable differential gene expression analysis for all samples (Fig. 2B). Moreover, both approaches yielded similar proportions of sequencing reads across the CDS regions, demonstrating minimal depletion method-induced bias (Fig. 3A). The average TPM of the housekeeping genes showed a conserved trend across different transcripts with DepStep yielding slightly higher TPM in comparison to commercial kit (Fig. 3B). This difference can be due to the transcriptional state of the input samples or a slight off-target potential of the commercial kit, although this cannot be verified as no list of target sequences is provided by the manufacturer.

Transcriptomic studies have become increasingly affordable as sequencing costs have plummeted, making such analyses more accessible to a broader number of researchers. However, RNA sequencing for a non-model prokaryotic organism remains expensive, owing to custom solutions needed for rRNA depletion. As we set out to develop a rRNA depletion method that would allow us to perform transcriptomic analysis on *S. glacialimarina*, we evaluated multiple commercial depletion kits of which only one kit yielded rRNA depletion. Alternatively, custom-made kits can be requested from commercial manufacturers, but such options are often prohibitively expensive for most research laboratories. Furthermore, our cost analysis of DepStep revealed that a significant portion of the cost is attributed to the commercially synthesized biotinylated probes. This expense can be minimized by performing in-house biotinylation at the 3’-end of the primer oligos (Flickinger et al. 1992). Nevertheless, even when accounting for the cost of commercially sourced biotinylated probes, DepStep reduced the cost-per-sample by a threefold compared to the commercial kit. Therefore, if cost is a major limiting factor and an exhaustive search for an optimal depletion method is to be avoided, then an in-house method using biotinylated beads, such as DepStep, presents a custom solution. In conclusion, by integrating key experimental design parameters, such as probe concentration, coverage, and inter-probe spacing, DepStep provides a consolidated workflow that simplifies the probe-based depletion strategy into a streamlined, cost-effective, single-step solution for gene expression analysis in non-model organisms.

## MATERIALS AND METHODS

### Bacterial strain

The bacterial strain used in this study is *Shewanella glacialimarina* TZS-4_T_ (DSM 115441, HAMBI 3773) and it was cultured as previously described (Qasim et al. 2021; Lampi et al. 2023). Briefly, *S. glacialimarina* was grown on solid media consisting of rich 25% w/v marine broth [abbreviated as rMB; 7.5 g peptone (Sigma-Aldrich), 1.5 g yeast extract (Fisher Bioreagents) and 9.35 g Marine broth (BD-Difco) in 1 L of deionized H_2_O] and agar (15 g/L) at 15 °C until colonies appear. For liquid culture, a 48 h overnight starter culture was grown from a single colony and used to inoculate 50 ml of 25% rMB to an OD_600_= 0.2. All the cultures were grown at 15 °C with constant aeration at 200 rpm. The cultures were grown until the desired OD_600_ and then the cells were harvested by centrifugation.

### Probe design for DepStep rRNA depletion

The rRNA sequences from 23S, 16S and 5S were aligned using MAFFT to generate a consensus sequence while replacing the variable region with “N” (Katoh 2002). The consensus sequence was then used to generate targeted antisense probes using default parameters in the oligostan.R script (Tsanov et al. 2016). The default parameters include minimum length = 26, maximum length = 32, ΔG_min_ = -28 kcal/mol, ΔG_max_ = -36 kcal/mol. The probes were also screened to remove repetitive nucleotide sequences longer than 4-5 nucleotides. The GC content of the probe was kept between 40-60%. Probes with a hairpin structure >-3 kcal/mol and homodimer or heterodimer formation capacity >-10 kcal/mol were removed from the candidate list. Furthermore, assuming a 300 mM salt concentration, the melting temperature (T_m_) of the probes was >70 °C, estimated based on the formula described by Sambrook (Sambrook 1989). The probe candidates were checked for off-target binding using BLAST (https://www.ncbi.nlm.nih.gov/) and BURST v0.99.8 (Al-Ghalith and Knights 2020). For BLAST, the reference transcripts database was aligned with the candidate probes using BLASTn with the following settings: parameters threshold = 0.05, match score = 2, mismatch score = -3, word size = 1, gap cost = 5, gap extension cost = 2. Probes with 8 or more sequence matches were considered as potential off-targets and removed from the list. The complete workflow is outlined in the Supplemental Material. The final set of antisense rRNA depletion probes, compiled based on the aforementioned criteria, were ordered from Metabion and dissolved to a final concentration of 100 µM in a resuspension buffer (10 mM Tris-HCl pH 8, 0.1 mM EDTA). Please refer to Supplemental Table 1 for complete details on the probes.

### Total RNA extraction from *Shewanella glacialimarina* TZS-4_T_

Total RNA extraction was performed using acidic phenol:1-bromo-3-chloropropane (BCP) as previously described (Qasim et al. 2021). Briefly, cells were grown in 50 ml of 25% rMB until it reached OD_600_ of 0.8. The cells were harvested at 3200 × *g*, 4 °C for 10 min. The harvested cells were resuspended in 4 ml of 0.9% NaCl solution then 4 ml acidic phenol pH 5.3 (Sigma-Aldrich Cat. No. P4682) 1 ml Trizol and 800 µl BCP (Acros Organics Cat. No. 106860010). Cells in the suspension were lysed by vortexing for 10 min in the presence of glass beads. The lysate was centrifuged for 10 min at room temperature (RT), the aqueous phase was collected and twice subjected to a re-extraction step with a phenol:BCP (5:1). The cleared supernatant containing RNA was then precipitated from the lysate by adding 2.5× vol. 99.6% EtOH and by incubating at -20 °C overnight. Total RNA was pelleted by centrifugation at 10,000 × *g* for 20 min at 4 °C. The RNA pellets were washed twice with 80% EtOH followed by air drying and resuspension in RNase-free ddH_2_O. The RNA quality was confirmed by agarose gel electrophoresis analysis [2% w/v agarose, Tris-borate EDTA (TBE) gel containing 4 µl/100 ml gel of Midori Green (Nippon Genetics Cat. No. MG04)]. Imaging was performed using a ChemiDoc MP imaging system (Bio-Rad).

### rRNA removal from total RNA using DepStep

Biotinylated probes targeting rRNA were mixed in equimolar amounts and a 100 pmol/µl probe mix solution was prepared. For rRNA depletion, 1 µg of *S. glacialimarina* total RNA was used with different amount of rRNA probe mix (5-300 pmol). The hybridization reaction was performed in 2.5 µl of 10× Ribohyb buffer (20× SSC, 0.1% Tween-20) (Thompson et al. 2020), 1 µl of RNase inhibitor (Promega Cat. No. N2615) adjusted with ddH_2_O to a final volume of 25 µl. This hybridization reaction was incubated for 5 min on a ThermoMixer C (Eppendorf) at 70 °C and mixed with an equal volume of Dynabeads^™^ MyOne^™^ Streptavidin C1 (Invitrogen Cat. No. 65002) which was pre-washed according to manufacturer’s instructions. The samples with the beads were incubated on a ThermoMixer C (Eppendorf) for 15-20 min at 50 °C with gentle shaking at 1250 rpm. After the biotin-streptavidin reaction, the beads were collected by placing the tubes in a magnetic stand for 5 min. The cleared supernatant was transferred into a new tube and cleaned up using RNA Clean and concentrator kit (ZYMO-RESEARCH Cat. No. R1015). The RNA samples were eluted from the Zymo-Spin™ IC Column (ZYMO-RESEARCH Cat. No. C1004-50) using 10 µl of RNase-free ddH_2_O water. The control sample did not contain probes for the hybridization reaction which was then followed by bead binding and clean-up. The RNA samples were loaded onto 6% urea polyacrylamide gel and visualized on Gel-doc XR (Bio-Rad) apparatus.

### rRNA removal from total RNA using a commercial depletion kit

RiboCop rRNA depletion kit (Lexogen Cat. No. 126) was used to benchmark DepStep. The rRNA-depleted samples were prepared according to manufacturer’s instructions. For depletion, 1 µg of *S. glacialimarina* total RNA was diluted to 26 µl of RNase-free water and 4 µl of hybridization solution (HS) was added. Next, 5 µl of probe mix for gram-negative bacteria (META, G-) was added and thoroughly mixed. The samples were denatured on a ThermoMixer C (Eppendorf) at 75 °C for 5 min with gentle shaking at 1250 rpm. After denaturing the temperature was dropped to 60 °C and further incubated for 30 min. The depletion beads (DB) were washed and resuspended in depletion solution (DS) and 30 µl was added to the RNA samples. The samples were mixed by pipetting up and down 8 times and placed on the ThermoMixer C (Eppendorf) at 60 °C for 15 min at 1250 rpm. After gently spinning down, the samples were placed on a magnetic stand for 5 min and the supernatant containing rRNA-depleted RNA was collected.

The rRNA-depleted samples were further purified using magnetic beads. A mixture of purification bead (PB) and purification solution (PS), 24 µl and 108 µl respectively was added to the rRNA-depleted sample and incubated for 5 min at room temperature. The sample tubes were placed on a magnetic stand for 10 min and the supernatant was discarded. The beads were washed with 120 µl of 80% EtOH without removing the tubes from the magnetic stand. The washes were repeated twice, and the beads were left to dry for 5 min. The RNA was eluted from the beads by 12 µl of elution buffer (EB) and transferred into a fresh tube. Samples were stored at - 80 °C until library preparation.

### RNA quality control

The RNA samples were analysed using an Agilent 4150 TapeStation automated electrophoresis device (Agilent Technologies). The sample buffer was equilibrated at room temperature for 30 min and 5 µl of sample buffer was mixed with 1 µl of RNA. The sample was vortexed for 1 min and spun down another minute, followed by heating at 72 °C for 3 min and cooling at 4 °C for 1 min on a VWR XT96 gradient thermal cycler (Avantor). Next, the sample was spun down for 1 min and the tube strip was loaded into the Agilent Tapestation. The RNA screen tape ladder (Agilent Technologies; part no: 5067-5578) was loaded into the first tube and prepared similarly as the RNA sample. Following electrophoresis, the results were visualized and analysed using the Agilent TapeStation Analysis software v4.1 (Agilent Technologies).

### RNA sequencing and data analysis

The DepStep samples along with commercial rRNA-depleted samples and total untreated RNA (NT = no treatment) were used to prepare RNA sequencing libraries. RNA sequencing libraries were prepared using CORALL RNA-Seq V2 Library Prep Kit with UDI 12 nt Set A1 (Lexogen Cat. No. 171.96). For total untreated RNA (NT), 1 µg of RNA was used for preparing its sequencing library. The sequencing libraries were prepared according to the manufacturer’s instruction. The sequencing libraries were amplified using a VWR XT96 gradient thermal cycler (Avantor) and the cycle number was estimated using quantitative RT-PCR using primers from the CORALL kit. The pooled libraries were sequenced on an Illumina NextSeq2000 with single-end sequencing for 100 cycles at the Vienna BioCenter Core Facilities (VBCF). The raw sequencing reads were demultiplexed and converted to fastq format using bcl2fastq. The unique molecular identifiers (UMIs) were extracted from the reads using UMI-tools command “umi_tools extract” (Smith et al. 2017). The adapters were removed using trimmomatic 3.39 with TruSeq3-SE.fa file and the adapter clipping parameters set to 2:30:10 (i.e. seed mismatched: palindrome clip: simple clip threshold). The trimmed reads were aligned back to *S. glacialimarina* reference using burrows-wheeler aligner (BWA-mem) and the aligned reads were sorted according to coordinates using samtools (Li et al. 2009). The bam files were indexed with samtools and then deduplication script was executed using UMI-tools command named “umi_tools dedup” (Smith et al. 2017). The reads aligning to each rRNA were counted using samtools. The sequencing reads (FASTQ format) generated in this study was deposited to European Nucleotide Archive (ENA) under the project accession number PRJEB92206.

### Northern blotting

For northern blot detection of 23S and 16S rRNA, 1 µg of total RNA was loaded on 1.5% formaldehyde-agarose gel in 1× MOPS buffer (41.86 g MOPS free acid, 4.1 g sodium acetate, 3.72 g disodium EDTA). The gel preparation and the sample run followed the protocol outlined in RNA: A laboratory manual 2011 (Rio et al. 2011). The RNA was transferred onto an Amersham Hybond-N+ nylon membrane (Cytiva) using capillary action with 20× SSC as transfer buffer and crosslinked in a UV crosslinker (Analytic Jena) for 5-10 s. The membrane was prehybridized using pre-hybridization buffer (3× Denhardt solution, 5× SSC, 0.1% SDS) at 45 °C for 2 h. DNA probes were denatured by heating at 70 °C for 3 min, then immediately cooled on ice. Hybridization was performed in pre-hybridization buffer supplemented with 200 µg/ml salmon sperm DNA (Invitrogen Cat. No. 15632-011) and incubated at 45 °C overnight. The following day, the membrane was washed with a series of stringency washing buffers and incubated in Str-HRP conjugate (ThermoFisher Scientific; Cat. No. 21130) (1:1500 dilution) in pre-hybridization buffer for 15 min at RT. Enhanced chemiluminescence kit solutions (ThermoFisher Scientific; Cat. No. 32209) were mixed in a 1:1 ratio and applied to the membrane. After 1 min of incubation the membrane was placed in Gel-doc XR (Bio-Rad) and the detection time was set to 300 s (Sambrook, 2001).

For detecting 5S rRNA, total RNA (1 µg) was loaded onto 15% urea polyacrylamide gel in 0.5× TBE and visualized on a Gel-doc XR (Bio-Rad) apparatus. The RNA was then transferred onto an Amersham Hybond-N+ nylon membrane (Cytiva) using Trans-Blot SD semi-dry transfer cell (Bio-Rad Cat. No. 1703957). The polyacrylamide gel, nylon membrane and Whatman paper were presoaked in 0.5× SSC and the sandwich was assembled with Whatman paper on the outer side, followed by the gel and the membrane inside of the sandwich. The transfer was done for 30 min at 115 mA and then nylon membrane was crosslinked afterwards in a UV crosslinker (Analytic Jena) for 5-10 s. The membrane pre-hybridization, probing, washing, and chemiluminescence detection was performed as mentioned above.

## ACKNOWLEDGEMENTS

The authors thank Salla Kalaniemi and Sari Korhonen for their valuable technical assistance. The authors wish to acknowledge CSC - IT Center for Science, Finland for computational resources and the Next Generation Sequencing Facility at Vienna BioCenter Core Facilities (VBCF), member of the Vienna BioCenter (VBC), Austria, for NGS services. We also extend our gratitude to all members of the RNAcious Laboratory for their insightful feedback and supportive discussions. This research was funded by the Research Council of Finland (Academy Project, grant number 354906; to L.P.S.) and the Novo Nordisk Foundation (Emerging Investigator in Biotechnology-Based Synthesis and Production, grant number NNF19OC0054454; to L.P.S.). M.S.Q. is a fellow of the Doctoral Programme in Microbiology and Biotechnology, University of Helsinki.

## Notes

### Competing Interest Statement

The authors have declared no competing interest.

